# 3D-equivariant graph neural networks for protein model quality assessment

**DOI:** 10.1101/2022.04.12.488060

**Authors:** Chen Chen, Xiao Chen, Alex Morehead, Tianqi Wu, Jianlin Cheng

## Abstract

**Motivation:** Quality assessment of predicted protein tertiary structure models plays an important role in ranking and using them. With the recent development of deep learning end-to-end protein structure prediction techniques of generating highly confident tertiary structures for most proteins, it is important to explore corresponding quality assessment strategies to evaluate and select the structural models predicted by them since these models have better quality and different properties than the models predicted by traditional tertiary structure prediction methods.

**Results:** We develop EnQA, a novel graph-based 3D-equivariant neural network method that is equivariant to rotation and translation of 3D objects to estimate the accuracy of protein structural models by leveraging the structural features acquired from the state-of-the-art tertiary structure prediction method - AlphaFold2. We train and test the method on both traditional model datasets (e.g., the datasets of the Critical Assessment of Techniques for Protein Structure Prediction (CASP)) and a new dataset of high-quality structural models predicted only by AlphaFold2 for the proteins whose experimental structures were released recently. Our approach achieves state-of-the-art performance on protein structural models predicted by both traditional protein structure prediction methods and the latest end-to-end deep learning method - AlphaFold2. It performs even better than the model quality assessment scores provided by AlphaFold2 itself. The results illustrate the 3D-equivariant graph neural network is a promising approach to the evaluation of protein structural models. AlphaFold2 features are important for improving protein model quality assessment and are complimentary with the geometric property features extracted from structural models.

**Availability:** The source code is available at https://github.com/BioinfoMachineLearning/EnQA.

**Supplementary information:** Supplementary data are available.

## 1 Introduction

Predicting the structures of proteins from their sequences is crucial for understanding their roles in various biological processes. Various computational methods have been developed to predict protein structure from sequence information (Arnold, et al., 2006; Baek, et al., 2021; Hou, J, et al., 2019; Jumper, et al., 2021; Senior, et al., 2020; Xu, 2019; Yang, et al., 2020). However, some predicted structures are still far from the true structure, especially for some proteins lacking critical information such as homologous structural templates or residue-residue co-evolution information in their multiple sequence alignments. Besides, many computational methods produce multiple outputs for one input sequence. Thus, it is important to estimate the accuracy of the predicted tertiary structural models, i.e., their similarity or discrepancy with the native but unknown structure. Such estimation can help select the best models from the predicted candidates and identify erroneous regions in the models for further refinement.

Many methods for model quality assessment (QA) have been developed. For example, PCONS (Wallner, et al., 2007) and ModFOLDclustQ (McGuffin, et al., 2010) use the comparison between 3D models to evaluate their quality. VoroMQA (Olechnovic and Venclovas, 2017) computes confidence scores based on the statistical potential of the frequencies of observed atom contacts. SBROD (Karasikov, et al., 2018) uses a smooth orientation-dependent scoring function with a ridge regression model.

Deep learning-based QA methods have been reported. DeepQA (Cao, et al., 2016) uses a deep belief network and different agreement metrics. ProQ4 (Hurtado, et al., 2018) uses the partial entropy of the sequence characteristics with a Siamese network configuration. GraphQA (Baldassarre, et al., 2021) tackles the QA protein with graph convolutional networks based on geometric invariance modeling. Ornate (Pagès, et al., 2019) and DeepAccNet (Hiranuma, et al., 2021) are based on voxelized spatial information of the predicted models and 2D/3D convolution networks. DeepAccNet is one of the best-performing methods in the QA category of the CASP14 competition (Kwon, et al., 2021).

The pioneering development of the end-to-end deep learning method for protein structure prediction - AlphaFold2 (Jumper, et al., 2021) that generated highly confident 3D structures for most protein targets in CASP14 as well as the recent release of a similar approach - RoseTTAFold (Baek, et al., 2021) presents notable improvements in structure prediction and brings new challenges for the model quality assessment task because traditional QA methods developed for evaluating structural models predicted by traditional methods may not work well for the models predicted by the new methods such as AlphaFold2(Kwon, et al., 2021). Since the software of the end-to-end approach, such as AlphaFold2 has been publicly released and is becoming the primary tool for tertiary structure prediction, it is important to develop corresponding quality assessment methods to evaluate their models. Furthermore, since AlphaFold2 generates structural models with a self-reported per-residue local distance difference test (lDDT) (Mariani, et al., 2013) quality score, new QA methods should outperform (1) the consensus evaluation of a predicted model by comparing it with the reference models predicted by AlphaFold2 and (2) the self-reported per-residue lDDT score for models provided by AlphaFold2. And it would be interesting to investigate if and how various information extracted from AlphaFold2 predictions can be used to enhance the quality assessment of 3D tertiary structural models. Finally, it is important to leverage the latest deep learning techniques of analyzing 3D objects.

The concept of rotation and translation equivariance in neural networks is useful for the analysis of rotation/translation-invariant properties of 2D and 3D objects in multiple domains, including 2D images (Cohen and Welling, 2016; Worrall, et al., 2017), quantum interactions (Schütt, et al., 2017), and 3D point clouds (Fuchs, et al., 2020; Satorras, et al., 2021; Thomas, et al., 2018). For equivariant networks, applying rotation and translation to the input results in a corresponding equivalent transformation to the output of the network. Invariance is a special case of equivariance, in which the same output is generated from the networks when such transformations are applied. Because the quality of a protein structural model is invariant to rotation and translation, it is desirable to use equivariant networks to predict model quality. As the locations of residues in a protein model can be represented as point clouds in 3D space, it is natural to represent a protein model as a graph, which can be equivariant to its rotation and translation. For example, the refinement step in RoseTTAFold (Baek, et al., 2021) uses an equivariant SE(3)-Transformer architecture to update the 3D coordinates. GNNRefine uses a graph convolution network with invariant features for protein model refinement.

In this work, we present EnQA, a 3D equivariant graph network architecture for protein model QA. We evaluate the performance of our method on three different test datasets: the CASP14 stage2 models, the models of the Continuous Automated Model EvaluatiOn (CAMEO), and a collection of AlphaFold2 predictions for recently released protein structures in the Protein Data Bank (PDB). EnQA achieves state-of-the-art performance on all three datasets. It can distinguish the high-quality structural models from other models and performs better than the self-reported lDDT score from AlphaFold2. To the best of our knowledge, our method is the first 3D-equivariant network approach to the problem of model quality assessment. It can effectively evaluate the quality of the models predicted by the current high-quality protein structure prediction methods such as AlphaFold2 that previous QA methods cannot.

## 2 Methods

In this section, we first describe the training and test datasets and data processing procedure. Then we define the input features to represent protein tertiary structures. Finally, we introduce the EnQA architecture and the implementation details.

### 2.1 Datasets

#### 2.1.1 CASP model quality assessment dataset

We use structural models from server predictions for CASP 8-14 protein targets (Stage two models if available) (Kwon, et al., 2021; Moult, et al., 1995) as one dataset, which can be downloaded from https://predic-tioncenter.org/download_area/. Models are first filtered by removing those with missing or inconsistent residues with respect to the corresponding experimental structure. The models from CASP 8-12 are used for training. The models from CASP13 are used to validate the neural network and select its hyperparameters. The models from CASP14 are used as the benchmark/test dataset. The details of the data preparation are available in Supplementary Notes 1.1. As a result, there are 109,318 models of 477 CASP8-12 targets used for training, 12,118 models of 82 CASP13 targets used for validation, and 9,501 models of 64 CASP14 targets for the final benchmark/test, respectively. The models in the CASP dataset were generated by traditional protein structure prediction methods during the CASP experiments between 2008 and 2020. The average quality of the models is much lower than the models predicted by the state-of-the-art method – AlphaFold2.

#### 2.1.2 AlphaFold2 model quality assessment dataset

To create a QA dataset containing protein structural models predicted by the latest end-to-end prediction method - AlphaFold2, we first collect protein structures in the AlphaFold Protein Structure Database (Tunyasuvunakool, et al., 2021) with corresponding experimental structures in Protein Data Bank (https://www.rcsb.org/) (Berman, et al., 2000; Burley, et al., 2020) released after the cutoff date (04/30/2018) of the structures on which AlphaFold2 was trained. In total, there are 4676 protein targets collected after filtering out identical ones. We divide these targets into training and test/benchmark sets with a 9:1 ratio. The AlphaFold2 models of the targets selected for training are combined with the training dataset from CASP 8-12 as the final training data. None of these targets has above 30% sequence identity threshold with any target in CASP14 benchmark dataset. The targets for the final AlphaFold2 benchmark/test dataset are selected by two criteria: (1) released after the start date of CASP14 (05/14/2020) and (2) having sequence identity <30% with any sequence in the training data, which is filtered by MMseqs2 (Steinegger and Söding, 2017). In total, 178 test targets are selected for the AlphaFold2 benchmark/test data after filtering.

For each of these targets above, we generate 5 AlphaFold2 models using the model preset of “casp14”, restricting templates only to structures available before CASP14 (i.e., max_template_date = “2020-05-14”) to make sure the AlphaFold2 models of the targets are generated with only the information available before their experimental structures were released. The AlphaFold2 models generate for the training targets are added into the training data. The AlphaFold2 models for the 178 test target form the final AlphaFold2 test/benchmark dataset. The details of generating AlphaFold2 models are available in Supplementary Notes 1.2.

#### 2.1.3 CAMEO model quality assessment dataset

To create an additional benchmark dataset, we use the recent models from Continuous Automated Model EvaluatiOn (CAMEO) (Robin, et al., 2021). We downloaded the protein structural models registered between 9/04/2021 to 11/27/2021, which include predictions from the latest predictors from different groups, such as RoseTTAFold (Baek, et al., 2021). Models are filtered by removing submissions contains only partial sequence of the corresponding target. In total, 38 targets with 945 structural models are selected for benchmarking. The preprocessing procedure for the CAMEO dataset is described in Supplementary Notes 1.3.

### 2.2 Features

We use a graph to represent a protein structural model, which contains node features and edge features. The 1D node feature has a shape (L, d), and the 2D edge feature has a shape (L, L, d) in which L is the number of residues in the model and d is the number of dimensions. The node feature describes the information of each residue, while the edge feature describes the information for each pair of residues. We briefly describe each type of features below.

#### 2.2.1 Node features

We use the 20-number one-hot representation to encode 20 amino-acid types of each residue. Following the spherical convolutions on molecular graphs (Igashov, et al., 2021), we use three types of features to characterize the *geometric property* for each residue: the solvent-accessible surface area, the size of Voronoi cell (Olechnovič and Venclovas, 2014), and the shortest topological distance to nearby solvent-accessible residues (also known as “buriedness”). In addition, we leverage the information from AlphaFold2 predictions made for the protein sequence of each model to generate the quality features for the model. AlphaFold2 predictions used for feature generation are made with the template database curated before the release date of the experimental structure of any target in the CASP14, CAMEO and AlphaFold2 test datasets. The lDDT score of each residue in a structural model to be evaluated with respect to an AlphaFold2 prediction for the same target (called a reference model) is used as a feature for the residue. The AlphaFold2 self-reported lDDT score for each residue in the reference model is also used as a feature measuring the confidence of the reference model. Here five AlphaFold2 reference models are used for generating features for the structural models of each target, 10 lDDT features are generated for each residue in each structural model. The final shape of the node features for each residue is (L, 33).

#### 2.2.2 Edge distance features

We first extract the logits from the distogram representation of the Alphafold2 predictions for a protein target, which represents the probability of the beta carbon (Cb) distance between two residues falling into predefined 64 distance bins, which has a shape (L, L, 64). From the 64-bin distogram, we then compute the probability of the distance error between two residues in a structural model falling into the 9 distance bins defined by lDDT as follows.

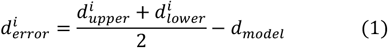

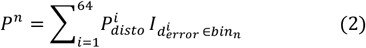

Where 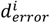is the distance error (difference) between the AlphaFold2 predicted distance and an input model for the *i-th* distance bin of Al- and phaFold2 and 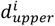 and 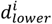 are the upper and lower bound of the *i-th* bin of the distogram, respectively. *d*_*model*_ is the distance between any two residues in the input model. *P*^*n*^ is the probability of the distance error between two residues falling into the *n-th* distance bin defined by lDDT (Mariani, et al., 2013). 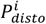 is the softmax-normalized probability of the i*-th* distance bin from AlphaFold2 distogram. *I* is an indicator function which equals 1 if 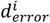falls into the range of the *n-th* bin defined by lDDT and 0 otherwise. The details of generating the pairwise distance error features of a model with respect to the distogram prediction of AlphaFold2 are available in Supplementary Notes 2.1. Since we use 5 AlphaFold2 distogram predictions for each target and 9 distance bins according to the definition of lDDT, this results in the pairwise edge features with a shape (L, L, 45) for each pair of residues in a structural model. We also create additional binary contact maps by summing up all probabilities in AlphaFold2 distograms that fall into the bins with middle point ≤ 15Å. The final binary contact map is the average from all five AlphaFold2 predictions to produce an additional edge feature with a shape (L, L, 1).

### 2.2.3 Spherical graph embedding edge features

We generate rotation-invariant graph embeddings following the Spherical Graph Convolutions Network (Igashov, et al., 2021) to use spatial information as spatial edge features. We first build the local coordinate frame for each residue in a structural model. We define the normalized Ca–N vector as the x-axis, the unit vector on the C–Ca–N plane and orthogonal to the Ca–N vector as the y-axis. The direction of the y-axis is determined by the one that has a positive dot product with the Ca–C vector. Naturally, the z-axis is the cross-product of x and y. We compute the spherical angles *θ* and *φ* of the vector between the Ca of each residue and that of any other residues with respect to this local spherical coordinate system. **Figure 1** illustrates the local spherical coordinate system used in this work.

**Figure 1.**
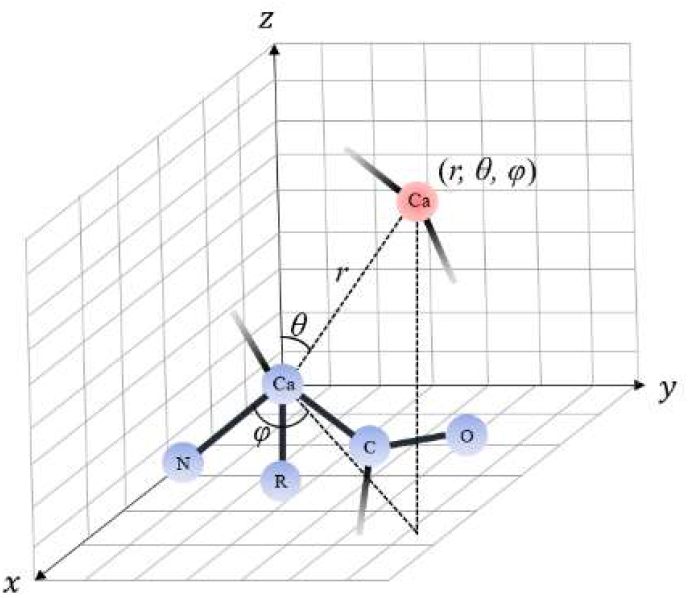
The illustration of the local spherical coordinate system. Different colors indicate atoms from different residues. Here *θ, φ* and *r* are spherical angles and the radial distance for the vector between the alpha carbons (Ca) of two residues (blue and red).

The spherical angles *θ* and *φ* are transformed into real spherical harmonics with the following formula:

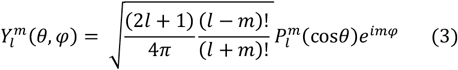

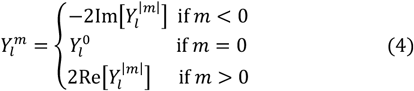

Here 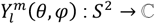 is a function defined on the surface of the unit sphere with degree *l* and order 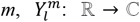 transform the complex spherical harmonics into their real forms. 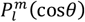 is the associated Legendre polynomials (Hobson, 1931). For spherical harmonics with degree *l* there are *2l+1* orders in total. We choose spherical harmonics with degrees from 0 to 4 in the graph embeddings, resulting in 25 orders for each pair of spherical angles *θ* and *φ*. The final graph embeddings have shapes (L, L, 25) and are concatenated with the pairwise edge distance features as model input. The structural information of the protein models is incorporated while preserving the rotation/translation invariance property by using such embeddings from the local spherical coordinate frame.

### 2.3 3D-equivariant model architecture

The overall architecture of our method is depicted in **Figure 2**. The processed 1D features (node features) are first processed with 1D convolutions to generate hidden node features. Then 2D features (both distance and graph embedding edge features) and the 2D tiling of the 1D hidden features are processed with a residual architecture with 5 blocks and 32 channels similar to the DeepAccNet (Hiranuma, et al., 2021). The goal is to predict an initial distance error as a classification task with 9 bins. The distance error is converted into an initial quality estimation using the binary contact map described in Section 2.2.2. The equation for the *n-th* residue in input with length L is the following:

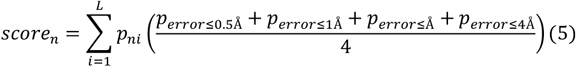

**Figure 2.**
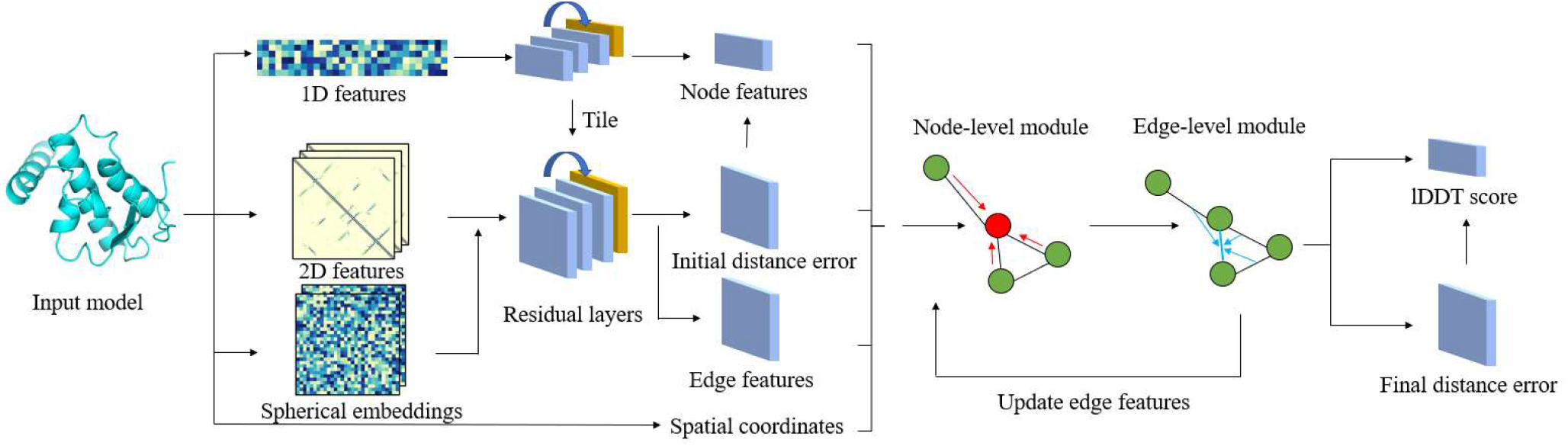
The illustration of the overall architecture of EnQA. The 1D/2D features from the input model are first converted into hidden node and edge features for the 3D-equivarant graph module. The spatial coordinates of Ca atoms of the residues are also used as an extra feature. The node and edge network modules update the graph features iteratively. In the end, the final per-residue lDDT score and distance errors of residue pairs are predicted from the updated node/edge features and spatial coordinates by the 3D-equivariant network.

Here *p*_*ni*_ is the probability of the beta carbon distance between *n-th* and *i-th* residue in the binary contact map. *p*_*error*_ is the sum of the probability of the multi-class error prediction from the residual layers below different distance cutoffs. This score is combined with the other 1D node features as the node features for the following 3D-equivariant graph network. The spatial coordinates of Ca atom of each residue from the input model are used as one additional feature, which is processed by the graph network in the 3D-equivariant manner and used to compute the final real value-based distance error. The input graph for the 3D-equivariant graph network is constructed by connecting any residue pairs with distance ≤ 15Å with an edge. The edge features for the graph network are the concatenation of the multi-class error prediction and a separate output of the residual layers for the pairs of the residues.

We use a variant of the E(n) Equivariant Graph Neural Networks (EGNN) (Satorras, et al., 2021) to process the node and edge features from the input graph and predict the final model quality score. Given a graph *G* = (*V, E*) with nodes *v*_*i*_ ∈ *V* and edges *e* _*ij*_ ∈ *E*. Our 3D-equvariant network has a node-level module and an edge-level module. In the node-level module, the hidden node features *h*_*i*_ ∈ ℝ and alpha carbon (Ca) coordinates *x*_*i*_ ∈ ℝ associated with each of the residues are considered. The equation of the EGNN layers is the following:

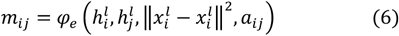

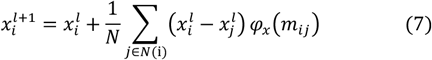

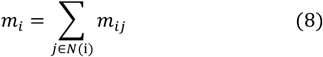

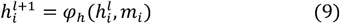

Here 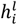 and 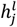 are the node features at layer *l, a*_*ij*_ is the edge feature,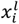 and 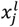 is the alpha carbon coordinates. *φ*_*e*_, *φ*_*x*_ and *φ*_*h*_ are multi-layer perceptron operations. *m*_*ij*_ and *m*_*i*_ are the intermediate messages for edges and nodes, respectively. The Ca coordinates are updated through each step so that its pairwise distance can reflect the distance map in the native PDB model, and can be used to compute the final real value-based distance error when subtracting the distance map from the initial coordinates of the model.

For the edge-level EGNN module, inspired by the Geometric Transformer (Morehead, et al., 2021), we use edges in the original graph as nodes, and define the new node features as the original edge features. Unlike the edges in the node-level module, we use the *k*-nearest neighbors approach to define the edges in the edge-level module with *k* set to 3 to accommodate the memory limit for edge-level graphs. The coordinates of the edges are the midpoint of two ends and are always determined by node coordinates rather than updates from the edge-level module. Finally, we use the distances between the midpoints as the new edge attributes. The whole architecture can be trained end-to-end from the input features to the final lDDT score prediction. In addition to the EGNN based graph layer, we also implemented a variant of the network by replacing the EGNN layers with a graph convolution network with kernels regularized by spherical harmonics functions as described in the SE(3)-Transformer (Fuchs, et al., 2020) for comparison.

We use 6 Nvidia Tesla V100 32G GPUs on the Summit supercomputer and Horovod/Pytorch to train the method. The batch size is set to 1 for each GPU, resulting in an effective batch size of 6. We use the stochastic gradient descent (SGD) optimizer with learning rate 1e-6, momentum 0.9 and weight decay 5e-5. We use the categorical cross-entropy as the loss function for initial distance error and the MSE loss for predicted lDDT scores as well as the final distance errors. The weight of the loss for predicted lDDT set to 5, while the weight of the other two errors is set to 1. We set the number of training epochs to 60 with early stopping when there are no improvements in validation loss for five consecutive epochs. Under our testing environment the deep learning model can handle proteins with length up to 850 residues. Structural models with sequence length longer than 850 are cropped into segments of length up to 800 and the final results are rebuilt with the concatenation of all the segments.

## 3 Results

### 3.1 Model quality assessment on CASP14 and CAMEO datasets

To compare the performance of EnQA with other state-of-the-art QA methods, we first evaluate it on the CASP14 stage 2 models (**Table 1**). We compare it with DeepAccNet (Hiranuma, et al., 2021), VoroMQA (Olechnovic and Venclovas, 2017) and ProQ4 (Hurtado, et al., 2018), which are all publicly available. We also use five AlphaFold models predicted for each CASP14 target as reference to evaluate the CASP14 stage 2 models. The average lDDT score between a CASP14 model and the five AlphaFold2 models is used as the predicted quality score of the model. This method is called AF2Consensus. The evaluation metrics used include residue and model-level mean squared error (MSE), mean absolute error (MAE) and Pearson Correlation Coefficient between the predicted lDDT scores and ground truth lDDT scores of the models. The per-residue metrics are first computed for each model and are then averaged across all models. Finally, the ranking losses in terms of lDDT and GDT-TS scores are used to evaluate the model ranking capability of the QA methods. The average of the predicted per-residue lDDT scores for each model is calculated as the predicted global quality score of the model. The predicted global quality scores for all the models for a target are used to rank them. The difference between the true GDT-TS (or true average lDDT score) of the best model and that of the top 1 ranked model of the target is the loss.

**Table 1.** The QA results on the CASP14 model dataset. Bold denotes the best result.

| Method | Per-residue |  |  | Per-model |  |  | Ranking loss |  |
| --- | --- | --- | --- | --- | --- | --- | --- | --- |
|  | MSE | MAE | Cor | MSE | MAE | Cor | IDDT | GDT-TS |
| AF2Consensus | 0.0057 | **0.0439** | 0.8596 | 0.0018 | 0.0244 | 0.9612 | 0.0092 | **0.0328** |
| DeepAccNet | 0.0254 | 0.1249 | 0.5725 | 0.0137 | 0.0945 | 0.7459 | 0.0444 | 0.0933 |
| VoroMQA | 0.0686 | 0.2115 | 0.3929 | 0.0466 | 0.1840 | 0.4620 | 0.0614 | 0.1175 |
| ProQ4 | 0.0296 | 0.1331 | 0.4493 | 0.0113 | 0.0806 | 0.7292 | 0.0570 | 0.1021 |
| EnQA-Full | **0.0049** | 0.0451 | **0.8676** | **0.0015** | **0.0227** | **0.9648** | **0.0088** | 0.0331 |
| EnQA-Reduced | 0.0477 | 0.1859 | 0.6470 | 0.0376 | 0.1790 | 0.8296 | 0.0408 | 0.0820 |
| EnQA-SE(3) | 0.0070 | 0.0607 | 0.7903 | **0.0015** | 0.0228 | 0.9611 | 0.0116 | 0.0323 |

Our method trained on the combination of CASP 8-12 models and AlphaFold2 models (EnQA-Full) outperforms all the other methods on both residue and model-level metrics (**Table 1**), except its per-residue MAE and ranking loss of GDT-TS is slightly worse than the consensus of AlphaFold2 (AF2Consensus). Compared with AF2Consensus, EnQA-Full achieves significant better per-residue MSE/Correlation, and per-model MSE/MAE, with p-value < 0.01 (paired sample t-test). EnQA-Full is performing better than AF2Consensus in terms of most metrics, demonstrating our 3D-equivariant QA method can add value on top of AlpahFold2 predictions in model quality assessment. EnQA-Full and AF2Consensus perform substantially better than the existing methods DeepAccNet, VoroMQA and ProQ4, indicating the importance of incorporating AlphaFold2 features into QA and the value of the 3D-equivariant architecture for QA. The variant of EnQA that uses the SE(3)-Transformer (EnQA-SE(3)) performs slightly worse than EnQA-Full, indicating the 3D-equivariant network (a variant of EGNN) in ENQA-Full may be slightly more effective. The model trained solely on AlphaFold2 models (EnQA-Reduced) yields the worse performance on the CASP14 test dataset than EnQA trained on both CASP8-12 and AphaFold2 models, which is expected since its training dataset contains only the models from AlphaFold2, which is not a good representative of the CASP14 server models generated by the traditional protein structure prediction methods. The quality of the former is generally much better than the latter.

We also evaluate all methods on the CAMEO dataset (**Table 2**). Similar to the results from the CASP14 benchmark dataset, EnQA-Full shows the best performances in all metrics, except the ranking loss, which falls behind AF2Consensus by a small margin. Compared with AF2Consensus, EnQA-Full achieves significantly better per-residue MSE/MAE/Correlation and per-model MSE/MAE, with p-value < 0.01 (paired sample t-test).

**Table 2.** The QA results on the CAMEO model dataset.

| Method | Per-residue |  |  | Per-model |  |  | Ranking loss |  |
| --- | --- | --- | --- | --- | --- | --- | --- | --- |
|  | MSE | MAE | Cor | MSE | MAE | Cor | IDDT | GDT-TS |
| AF2Consensus | 0.0084 | 0.0535 | 0.8529 | 0.0036 | 0.0353 | 0.9191 | **0.0054** | **0.0105** |
| DeepAccNet | 0.0245 | 0.1215 | 0.6636 | 0.0146 | 0.1006 | 0.7250 | 0.0144 | 0.0193 |
| VoroMQA | 0.1297 | 0.3175 | 0.4561 | 0.1094 | 0.3125 | 0.5512 | 0.0470 | 0.0537 |
| ProQ4 | 0.0684 | 0.2163 | 0.4498 | 0.0508 | 0.1961 | 0.5374 | 0.0656 | 0.0673 |
| EnQA-Full | **0.0061** | **0.0508** | **0.8602** | **0.0017** | **0.0272** | **0.9517** | 0.0068 | 0.0132 |
| EnQA-Reduced | 0.0265 | 0.1267 | 0.7250 | 0.0182 | 0.1159 | 0.8322 | 0.0177 | 0.0342 |
| EnQA-SE(3) | 0.0085 | 0.0681 | 0.7764 | 0.0021 | 0.0340 | 0.9335 | 0.0115 | 0.0190 |

### 3.2 Model quality assessment on AlphaFold2 dataset

To further assess the performance of our method on generally highquality models, we perform the evaluation on our AlphaFold2 test dataset (**Table 3**). We also include the self-reported lDDT scores of the models from AlphaFold2 as the baseline method for comparison (AF2-plddt). In this test, EnQA-Reduced, which trained solely on AlphaFold2 models, outperforms all other methods on all residue- and model-level metrics, indicating the importance of ensuring the consistency between the training models and test models. Its better performance than AF2-plddt shows that our method performs better in evaluating AlphaFold2 models than AlphaFold2’s own quality scores. Compared with AF2plddt, both EnQA-Reduced and EnQA-Full achieve significantly better per-residue MSE/MAE/Correlation and per-model MSE/MAE, with p-value < 0.01 (paired sample t-test). Furthermore, EnQA-full also outperforms all other methods except EnQA-Reduced in all metrics, including AF2-plddt, indicating combining AlphaFold2 models with traditional protein structural models for training the deep learning method can work well on both new AlphaFold2 test models and non-AlphaFold test models. All our three methods perform substantially better than the previous QA methods (DeepAccNet, VoroMQA, and ProQ4) on this dataset, clearly demonstrating the need of developing new QA methods for evaluating AlphaFold2 models.

**Table 3.** The QA results on the AlphaFold2 model dataset.

| Method | Per-residue |  |  | Per-model |  |  | Ranking loss |  |
| --- | --- | --- | --- | --- | --- | --- | --- | --- |
|  | MSE | MAE | Cor | MSE | MAE | Cor | IDDT | GDT-TS |
| AF2-plddt | 0.0232 | 0.1119 | 0.6651 | 0.0148 | 0.1011 | 0.7113 | 0.0052 | 0.0139 |
| AF2Consensus | 0.0417 | 0.1579 | 0.5549 | 0.0314 | 0.1562 | 0.6125 | 0.0098 | 0.0235 |
| DeepAccNet | 0.0394 | 0.1518 | 0.4907 | 0.0269 | 0.1426 | 0.5255 | 0.0086 | 0.0226 |
| VoroMQA | 0.1887 | 0.3899 | 0.3892 | 0.1644 | 0.3856 | 0.2386 | 0.0090 | 0.0233 |
| ProQ4 | 0.0983 | 0.2690 | 0.4111 | 0.0791 | 0.2565 | 0.2857 | 0.0103 | 0.0246 |
| EnQA-Full | 0.0132 | 0.0803 | 0.6994 | 0.0058 | 0.0533 | 0.7439 | 0.0049 | 0.0123 |
| EnQA-Reduced | **0.0118** | **0.0768** | **0.7090** | **0.0043** | **0.0455** | **0.7814** | **0.0046** | **0.0109** |
| EnQA-SE(3) | 0.0127 | 0.0840 | 0.6556 | 0.0045 | 0.0511 | 0.7540 | 0.0073 | 0.0178 |

### 3.3 Analysis of the performance on AlphaFold2 predicted models

We first examine the distribution of model quality of the models in the AlphaFold2 test dataset (**Figure 3**). The average true lDDT score for all models is 0.8034, with 79.82% above 0.7. The distribution of model quality of the CASP and CAMEO datasets are provided in **Figure S1** and **S2**. The results indicate that the structure models in the AlphaFold2 test dataset have much higher average quality than the CASP and CAMEO test datasets.

**Figure 3.**
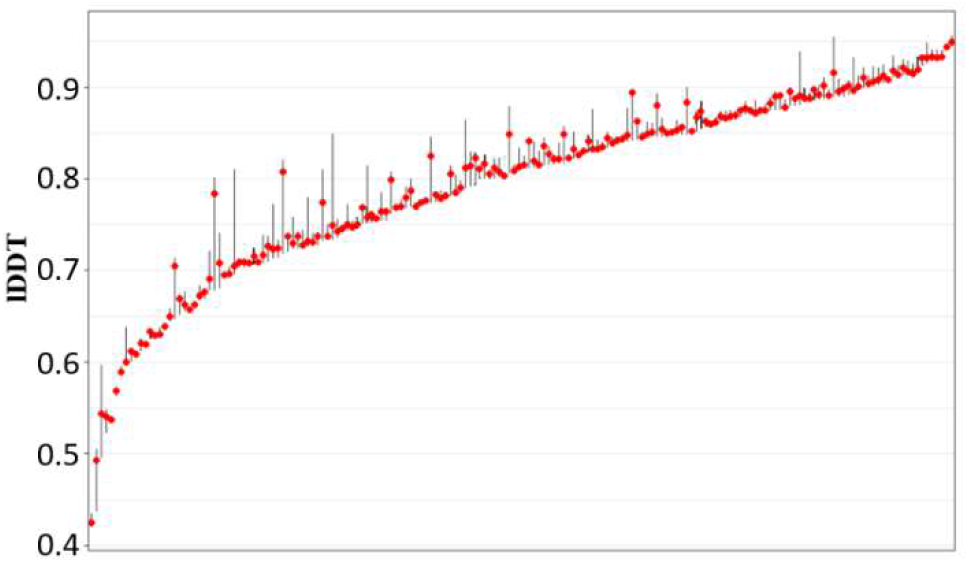
The distribution of lDDT scores of AlphaFold test models. X axis denotes the targets ordered by the mean lDDT of their models in increasing order. The red dots indicate the position of the median.

We further investigate the characteristics of the predictions of EnQA-Full and the self-reported lDDT score from AlphaFold2 predictions on the AlphaFold2 test models (**Figure 4**). The predicted scores of EnQA-Full have higher correlation with the true lDDT scores than AlphaFold2 self-reported quality scores. At both the residue and global-level, the AlphaFold2 reported score tends to overestimate the quality of the models more than EnQA-Full, which explains one improvement made by EnQA-Full.

**Figure 4.**
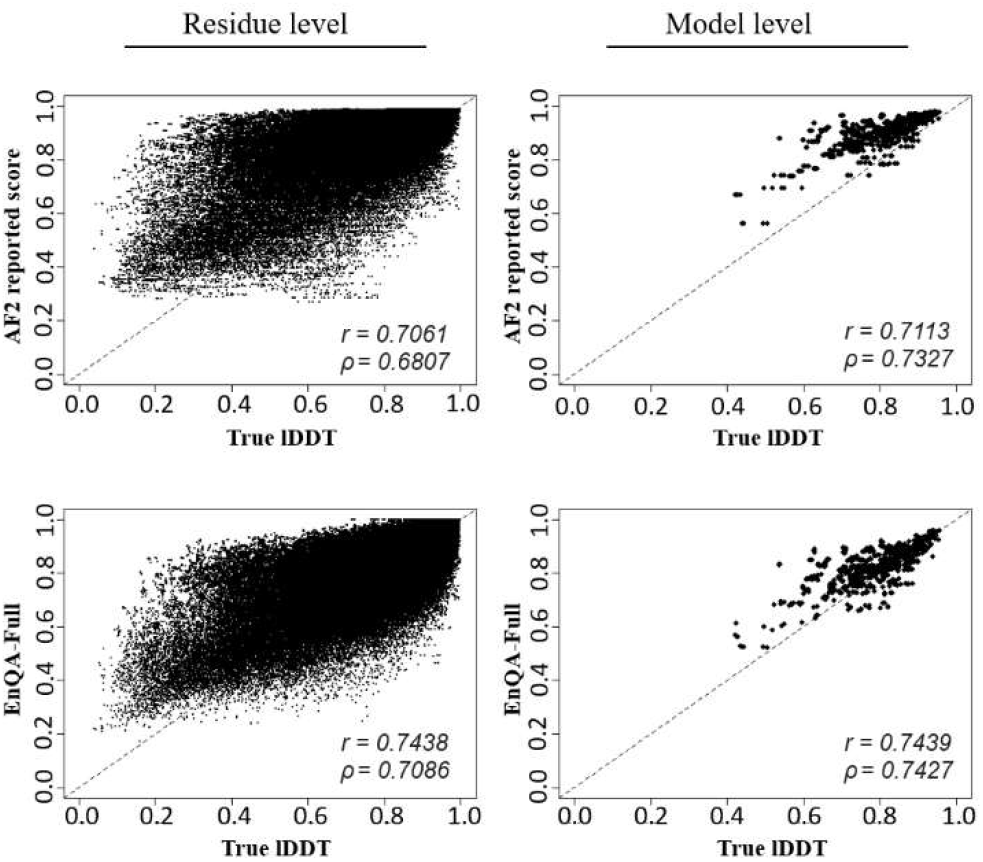
The comparison between the predicted and true lDDT scores for AlphaFold models. The residue-level correlation is computed for all residue at once, which is different from the average of the residue-level correlation in each model (used by Sections 3.1 and 3.2). *r*: Pearson Correlation Coefficient. *ρ*: Spearman Correlation Coefficient

In addition, we plot the true lDDT scores and the absolute error of the predicted scores on the AlphaFold2 test models (**Figure S3**). For both AlphaFold2 self-reported lDDT scores and EnQA predicted lDDT scores, the errors mainly come from overestimating the quality of the low-quality residues in the models.

### 3.4 Analysis of the impact of features

We examine the impact of different input features on the prediction performance of EnQA. We calculate the residue-level Pearson’s Correlation Coefficient between predicted lDDT score and true lDDT score when each type of feature is replaced with random number on CASP14 and AlphaFold2 test datasets (**Figure 5**). We use EnQA-Full as the baseline model and report the prediction performance when each feature is randomly permuted in its value range during prediction. A larger change of Pearson Correlation Coefficient indicates a higher impact. The lDDT score feature of a model with respect to AlphaFold reference models is the most important feature on both CASP14 and AlphaFold2 test datasets as its permutation causes the largest drop of the Pearson’s Correlation Coefficient. The geometric property features for each node, distograms, and the confidence score of AlphaFold also have a noticeable impact on the predictive capability. The similar trend can also be observed on the CAMEO dataset (**Figure S4**).

**Figure 5.**
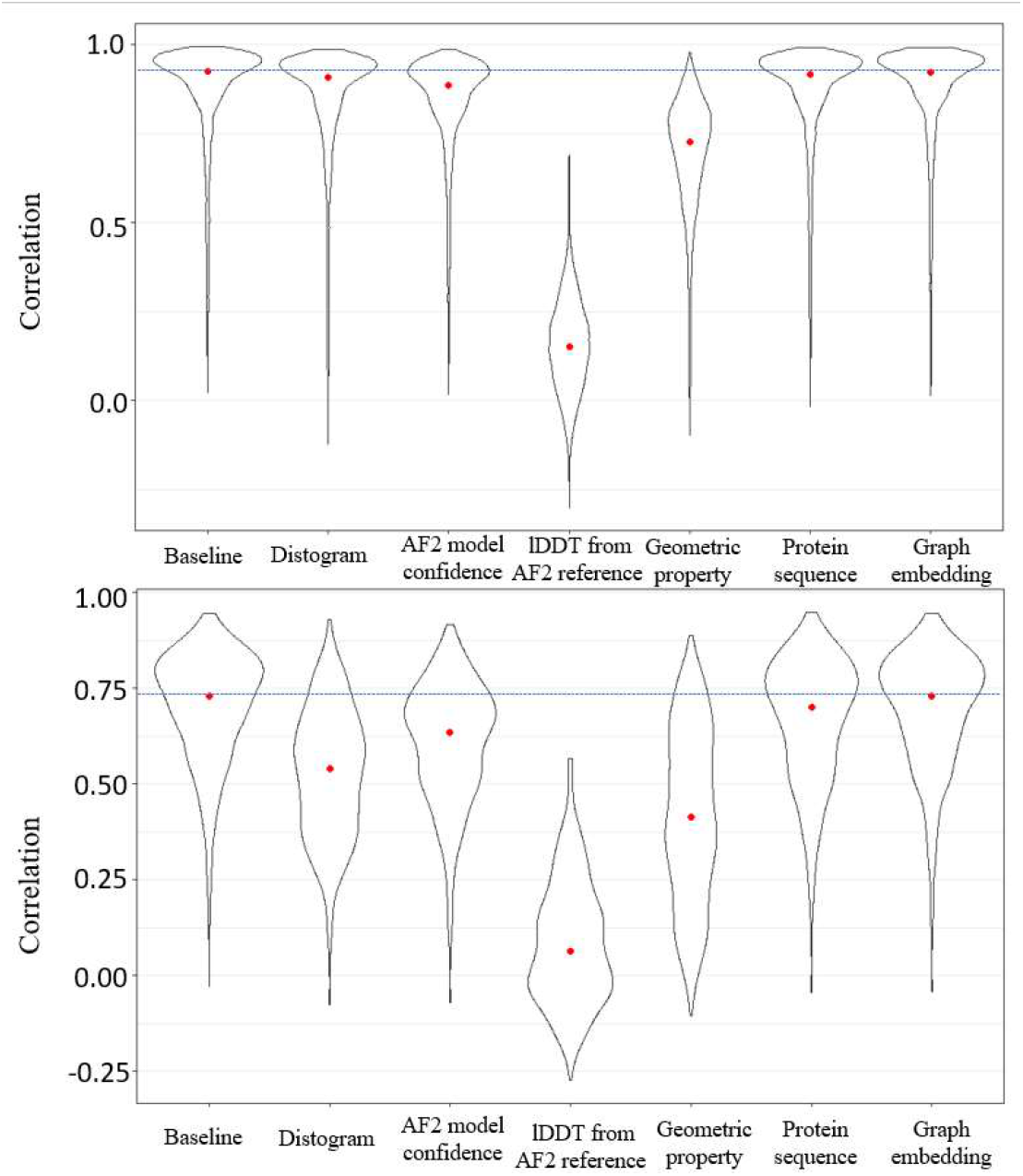
The comparison of residue-level Pearson’s Correlation Coefficient when different features are randomly permuted for model quality assessment. The red dots indicate the position of the median. Top - on the CASP14 test dataset; bottom - on AlphaFold2 test dataset.

Furthermore, we show that the geometric property feature is critical for those models on which EnQA-full makes large improvements over AF2Consesus. We pick the top 10% models for which it has the largest improvements in residue-level correlation over AF2Consensus and bottom 10% models for which it has the least improvement. In **Figure 6**, the importance of the geometric property feature (i.e., the change of the correlation) in top 10% models is significantly higher than the bottom 10% models (p value < 0.01 according to Mann-Whitney test), suggesting the orthogonal, synergistic effect of the geometric property feature and the features extracted from AlphaFold2 predictions.

**Figure 6.**
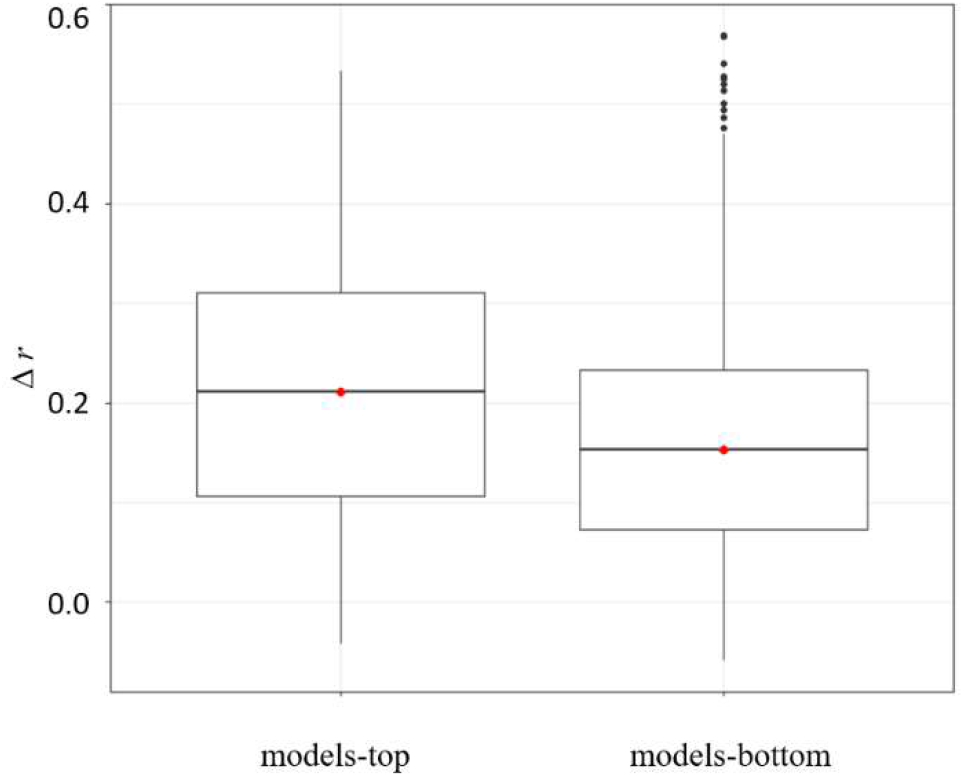
The comparison of the importance of the geometric property feature measured as the decrease in residue-level correlation for models that have most (models-top) and least (models-bottom) improvements in EnQA-Full over AF2Consensus. The change in the correlation in the former is higher than in the latter (*p* < 0.01, Mann-Whitney test).

## 4 Conclusion

In this paper, we introduce EnQA, a novel 3D-equivariant network method for protein quality assessment. Our approach utilizes both the geometric structural features of an input model and the features extracted from AlphaFold2 predictions. The network is developed as an equivariant framework with the node and edge features passing through the node and edge-level graph networks. Performed computational experiments on diverse structural model datasets prove EnQA achieves the state-of-the-art performance of protein quality assessment. More precisely, on both CASP14 and recent CAMEO protein structures, EnQA outperforms all other methods on most evaluation metrics, including using AlphaFold2 predictions as reference to evaluate models. Furthermore, our method performs better than the self-reported lDDT score of AlphaFold2 in evaluating high-quality AlphaFold2 models. On all the test datasets, EnQA performs substantially better than the previous QA methods, demonstrating the value of using 3D-equivarnant architecture and AlphaFold2-based features. Also, we show that the input features extracted from structural models have a complementary effect with the information extracted from AlphaFold2 predictions, especially for those models on which EnQA performs better.

To the best of our knowledge, the method is the first 3D-equivariant network approach to leveraging information from AlphaFold2 predictions to improve model quality assessment. It may be further expanded for protein model refinement by adding additional graph layers to update the coordinates of the nodes (residues) and other protein structure analysis tasks.

## Acknowledgements

This research used resources of the Oak Ridge Leadership Computing Facility, which is a DOE Office of Science User Facility supported under Contract DE-AC05-00OR22725.

## References

Arnold, K., et al. The SWISS-MODEL workspace: a web-based environment for protein structure homology modelling. Bioinformatics 2006;22(2):195–201.

Baek, M., et al. Accurate prediction of protein structures and interactions using a three-track neural network. Science 2021;373(6557):871–876.

Baldassarre, F., et al. GraphQA: protein model quality assessment using graph convolutional networks. Bioinformatics 2021;37(3):360–366.

Berman, H.M., et al. The Protein Data Bank. Nucleic Acids Res 2000;28(1):235–242.

Burley, S.K., et al. RCSB Protein Data Bank: powerful new tools for exploring 3D structures of biological macromolecules for basic and applied research and education in fundamental biology, biomedicine, biotechnology, bioengineering and energy sciences. Nucleic Acids Research 2020;49(D1):D437–D451.

Cao, R., et al. DeepQA: improving the estimation of single protein model quality with deep belief networks. BMC bioinformatics 2016;17(1):1–9.

Cohen, T. and Welling, M. Group equivariant convolutional networks. In, International conference on machine learning. PMLR; 2016. p. 2990–2999.

Fuchs, F.B., et al. Se (3)-transformers: 3d roto-translation equivariant attention networks. arXiv preprint arXiv:.10503 2020.

Hiranuma, N., et al. Improved protein structure refinement guided by deep learning based accuracy estimation. Nature Communications 2021;12(1):1340.

Hobson, E.W. The theory of spherical and ellipsoidal harmonics. CUP Archive; 1931.

Hou, J., Wu, T., Cao, R., & Cheng, J. Protein tertiary structure modeling driven by deep learning and contact distance prediction in CASP13. Proteins: Structure, Function, and Bioinformatics, 2019;87(12), 1165–1178

Hurtado, D.M., Uziela, K. and Elofsson, A. Deep transfer learning in the assessment of the quality of protein models. arXiv preprint arXiv:.06281 2018.

Igashov, I., Pavlichenko, N. and Grudinin, S. Spherical convolutions on molecular graphs for protein model quality assessment. Machine Learning: Science and Technology 2021;2(4):045005.

Jumper, J., et al. Highly accurate protein structure prediction with AlphaFold. Nature 2021;596(7873):583–589.

Karasikov, M., Pagès, G. and Grudinin, S. Smooth orientation-dependent scoring function for coarse-grained protein quality assessment. Bioinformatics 2018;35(16):2801–2808.

Kwon, S., et al. Assessment of protein model structure accuracy estimation in CASP14: Old and new challenges. Proteins 2021;89(12):1940–1948.

Mariani, V., et al. lDDT: a local superposition-free score for comparing protein structures and models using distance difference tests. Bioinformatics 2013;29(21):2722–2728.

McGuffin, L. J., & Roche, D. B. Rapid model quality assessment for protein structure predictions using the comparison of multiple models without structural alignments. Bioinformatics, 2010;26(2), 182–188

Morehead, A., Chen, C. and Cheng, J. Geometric Transformers for Protein Interface Contact Prediction. arXiv preprint 024232021.

Moult, J., et al. A large-scale experiment to assess protein structure prediction methods. Proteins 1995;23(3):ii–v.

Olechnovic, K. and Venclovas, C. VoroMQA: Assessment of protein structure quality using interatomic contact areas. Proteins 2017;85(6):1131–1145.

Olechnovič, K. and Venclovas, C. Voronota: A fast and reliable tool for computing the vertices of the Voronoi diagram of atomic balls. J Comput Chem 2014;35(8):672–681.

Pagès, G., Charmettant, B. and Grudinin, S. Protein model quality assessment using 3D oriented convolutional neural networks. Bioinformatics 2019;35(18):3313–3319.

Robin, X., et al. Continuous Automated Model EvaluatiOn (CAMEO)-Perspectives on the future of fully automated evaluation of structure prediction methods. Proteins 2021;89(12):1977–1986.

Satorras, V.G., Hoogeboom, E. and Welling, M. E (n) equivariant graph neural networks. arXiv preprint 098442021.

Schütt, K.T., et al. SchNet: a continuous-filter convolutional neural network for modeling quantum interactions. In, Proceedings of the 31st International Conference on Neural Information Processing Systems. Long Beach, California, USA: Curran Associates Inc.; 2017. p. 992–1002.

Senior, A.W., et al. Improved protein structure prediction using potentials from deep learning. Nature 2020;577(7792):706–710.

Steinegger, M. and Söding, J. MMseqs2 enables sensitive protein sequence searching for the analysis of massive data sets. Nature Biotechnology 2017;35(11):1026–1028.

Thomas, N., et al. Tensor field networks: Rotation-and translation-equivariant neural networks for 3d point clouds. arXiv preprint 082192018.

Tunyasuvunakool, K., et al. Highly accurate protein structure prediction for the human proteome. Nature 2021;596(7873):590–596.

Wallner, B., Larsson, P., & Elofsson, A. Pcons. net: protein structure prediction meta server. Nucleic acids research, 2007;35(suppl_2), W369–W374

Worrall, D.E., et al. Harmonic networks: Deep translation and rotation equivariance. In, Proceedings of the IEEE Conference on Computer Vision and Pattern Recognition. 2017. p. 5028–5037.

Xu, J. Distance-based protein folding powered by deep learning. Proceedings of the National Academy of Sciences 2019;116(34):16856–16865.

Yang, J., et al. Improved protein structure prediction using predicted interresidue orientations. Proceedings of the National Academy of Sciences 2020;117(3):1496–1503.

