## Supplementary for "3D-equivariant graph neural networks for protein model quality assessment"

### Supplementary Notes

#### 1. Selection of Datasets

##### 1.1 CASP14 benchmark dataset protocol

CASP14 target list from the QA results (68 in total)
https://predictioncenter.org/casp14/targetlist.cgi?view=regular&assis_type=all&view=all&field=t.release_date&order=ASC

Remove 6 targets ('T1048', 'T1072s1', 'T1062', 'T1070', 'T1080', 'T1077'
) that are not evaluated in global QA benchmark as is described in the official assessment (66 remaining) https://onlinelibrary.wiley.com/doi/full/10.1002/prot.26192

Remove two targets without publicly available native structure (64 remainings)
"T1085" "T1086"

"T1098 MESHI_SERVER_TS4" is excluded due to incomplete prediction compared to the reference structure (486/538 residues).

##### 1.2 AlphaFold2 dataset

We use 5 models from our AlphaFold predictions and the models from AlphaFold database for training, and for testing we use 5 models from our AlphaFold predictions. The configuration we used for running AlphaFold2 is setting the parameters to "full_dbs" and max_template_date is set to "2020-05-14".

For selecting targets for training and benchmark, we first sample 10% (468) of the count of all targets from those targets released after 05/14/2020. The rest are combined with the training datasets created using targets in CASP 8-12. Then we use MMseqs2 to filter out any targets that have sequence identity <30% with any sequence in the training data. Finally, the remaining 178 targets are used for further analysis in the benchmark.

##### 1.3 Targets from CAMEO dataset

We first filter the models by removing those predictions with inconsistent sequences with the corresponding reference structure. Targets are excluded for further analysis if less than three models are left after filtering.

#### 2. Feature generation

##### 2.1 Distance error

The segregation of the lDDT bins is defined as [-∞, -4.0, -2.0, -1.0, -0.5, 0.5, 1.0, 2.0, 4.0, +∞]
The segregation of AlphaFold distogram bins is defined as [2.000, 2.3125 , 2.625 , 2.9375 , 3.25 , 3.5625 , 3.875 ,4.1875 , 4.5 , 4.8125 , 5.125 , 5.4375 , 5.75 ,6.0625 ,

6.375 , 6.6875 , 7. , 7.3125 , 7.625 , 7.9375 , 8.25 , 8.5625 , 8.875 , 9.1875 , 9.5 , 9.812499, 10.125 , 10.4375 , 10.75 , 11.0625 , 11.375 , 11.6875 , 12. , 12.3125 , 12.625 , 12.9375 , 13.25 , 13.5625 , 13.875 , 14.1875 , 14.5 , 14.8125 , 15.125 , 15.4375 , 15.75 , 16.0625 , 16.375 , 16.6875 , 17.000 , 17.312498, 17.625 , 17.9375 , 18.25 , 18.5625 , 18.875 , 19.1875 , 19.5 , 19.8125 , 20.125 , 20.4375 , 20.75 , 21.0625 , 21.375 , 21.6875, 22.000 ]

For any residue pair of distance $d_{model}$ in the model, we define the distance error between the predicted Alphafold model and input model for the *i-th* distance bin of Alphafold as:

$$d_{error}^{i} = (d_{upper}^{i} + d_{lower}^{i})/2 - d_{model}$$

Here $d_{upper}^{i}$ and $d_{lower}^{i}$ are the upper and lower bound of the *i-th* bin of the distogram.

We then compute the probability of the distance error between two residues falling into the *n-th* distance bin defined by lDDT as:

$$P^{n} = \sum_{i=1}^{64} P_{disto}^{i} \, I_{d_{error}^{i} \in bin_{n}}$$

Here $P_{disto}^{i}$ is the Softmax-normalized probability of the i-*th* distance bin from Alphafold distogram. $I_{d_{error}^{i} \in bin_{n}}$ is an indicator function which equals 1 if $d_{error}^{i}$ falls into the *n-th* bin defined by lDDT and 0 otherwise. In practice, we set the first segregation of AlphaFold bins to 0.

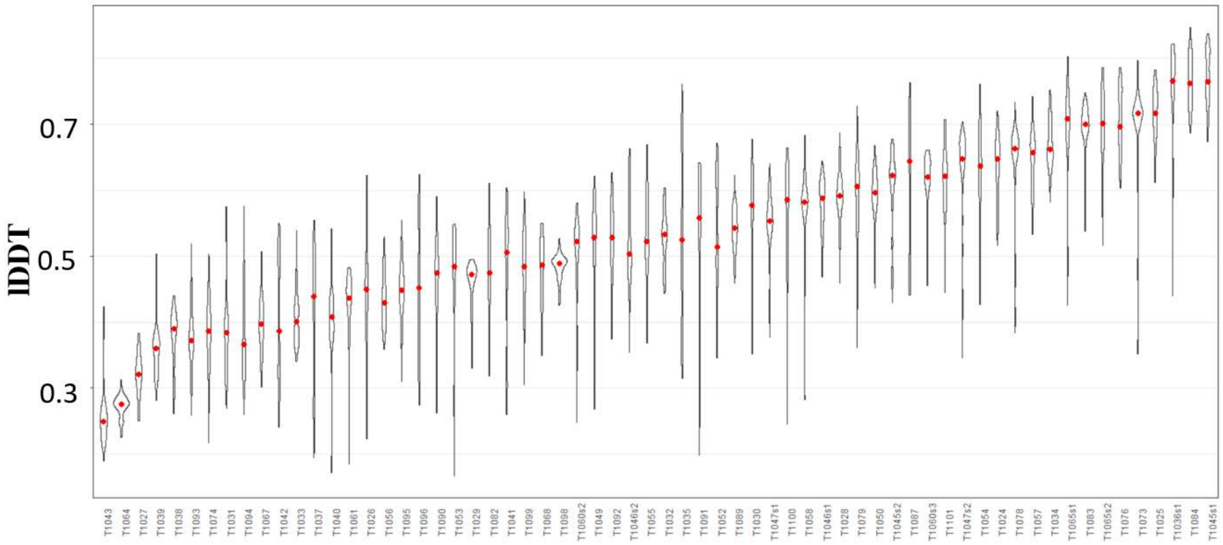


**Figure S1.** The distribution of lDDT score in the benchmark dataset for CASP14 models. The targets are ordered by mean lDDT. The red dots indicate the position of the median.

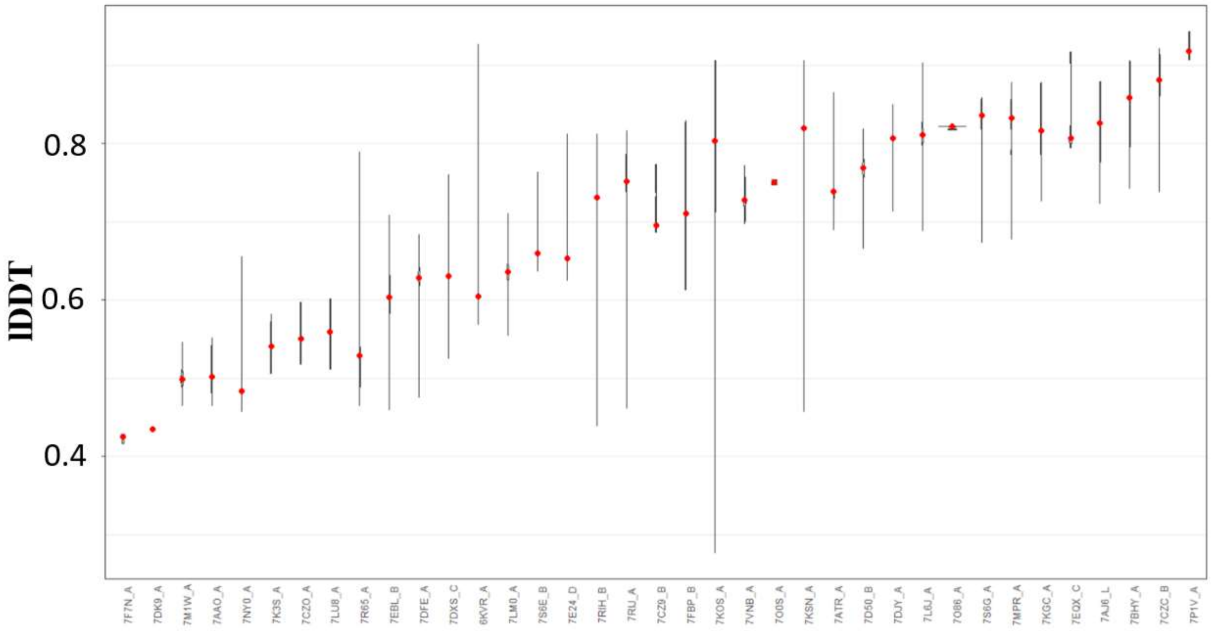


**Figure S2.** The distribution of lDDT score in the benchmark dataset for CAMEO models. The targets are ordered by mean lDDT. The red dots indicate the position of the median.

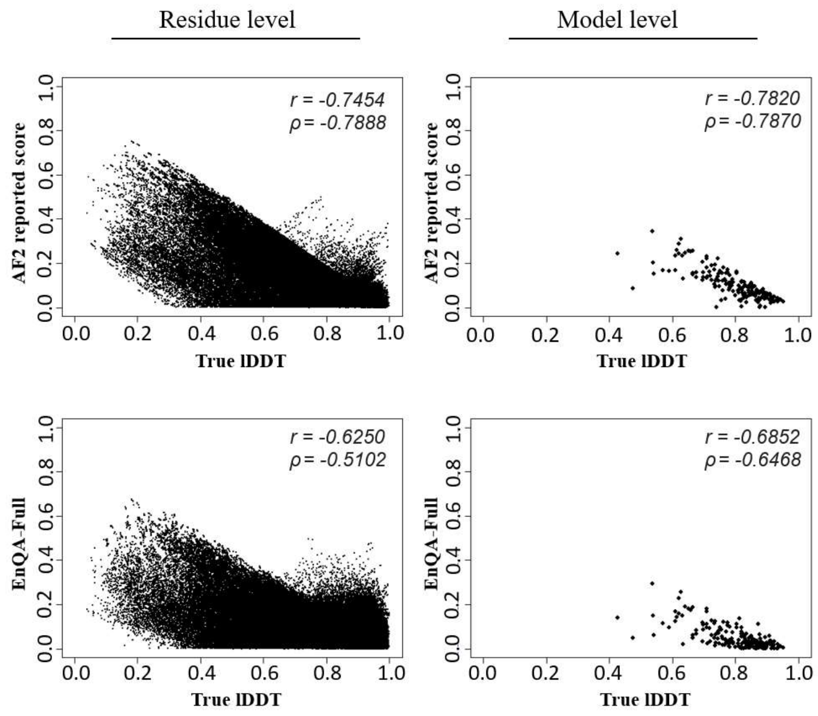


**Figure S3.** The comparison between the absolute error in prediction and true lDDT score for AlphaFold models.

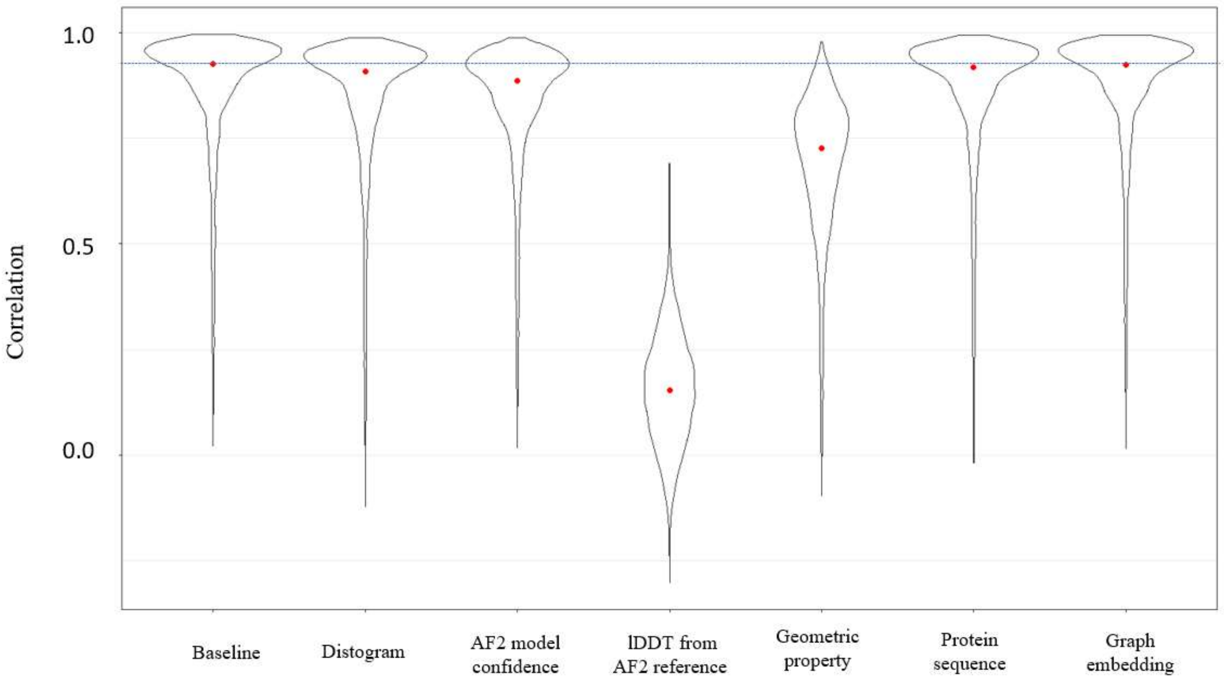


**Figure S4.** The comparison of residue-level Pearson Correlation Coefficient when different features are excluded for model quality assessment on CAMEO dataset. The red dots indicate the position of the median.
